# Dynamic Mechanical Cue Facilitate Collective Responses of Crowded Cell Population

**DOI:** 10.1101/2020.05.19.103275

**Authors:** Bingchen Che, Wei Zhao, Guangyin Jing, Jintao Bai, Ce Zhang

## Abstract

Collective cell behavior is essential for tissue growth, development and function, e.g. heartbeat^1^, immune responses^2^ and cerebral consciousness^3^. In recent years, studies on population cells uncover that collective behavior emerges in both inter- and intra-cellular activities, e.g. synchronized signal cascade^4^, and collective migration^5^. As the movement and shape transition of cells within the crowded environment of biological tissue can generate mechanical cues at the cell-cell interface, which may affect the signaling cascade^6,7^, we suspect that the inter- and intra-cellular collective behavior interplay with one another and cooperatively regulate life machinery. To verify our hypothesis, we study the collective responses of fibroblasts in a confluent cell monolayer (CCM). Our results demonstrate that cells in CCM show distinctive behavior as compared to the stand-alone (SA) cells, suggesting effect of inter-cellular interactions. Upon periodic TNF-α stimulation, collective behavior emerges simultaneously in NF-κB signaling cascade and nuclear shape fluctuations in CCM but not SA cells. We then model the inter-cellular interactions in CCM using a customized microfluidic device, and discover a feedback loop intrinsic to CCM, in which dynamic mechanical cues and mechano-signaling act as link connecting the inter- and intra-cellular collective activities. We found that mechano-signaling triggered by the dynamic mechanical cues causes collective nuclear shape fluctuation (NSF), which subsequently facilitates the collective behavior in NF-κB dynamics. Furthermore, our studies reveal that regardless of the input TNF-α periodicity, cellular responses of single fibroblasts are elevated when the dynamic mechanical cues synergize with the chemical inputs, and inhibited when there is phase-mismatching. We, therefore, postulate that besides the biological significance of mechano-signaling in regulating collective cell responses, the induction of dynamic mechanical cues to human body may be a potential therapeutic approach, allowing us to regulate the action of single cells to achieve optimal tissue performance.

## Introduction

To accomplish most biological functions (e.g. embryogenesis, tumorigenesis, and immune responses), population cells must act collectively, which allows the living tissue to undergo drastic morphological and behavioral transitions^5,8–13^. The collective behavior of population cells can be categorized into two groups: the inter- and intra-cellular activities. On one hand, the inter-cellular interaction, e.g. short-range cell-cell and cell-extracellular matrix (ECM) interactions lead to collective motion, transitions in cell and tissue shape^5,8,12^, which defines the biological tissue as active material. The intra-cellular activities (i.e. signaling cascade) of population cells, on the other hand, exhibit collective behavior when being exposed to dynamic chemical stimuli^4^. The mammalian immune response is a striking example^14^. Tay, et al. revealed that fibroblasts are capable of processing dynamic signals^15^. At a defined signal input frequency, the otherwise random signaling cascade of population cells become coordinated, leading to enhanced immune responses.

An important open question is whether inter- and intra-cellular collective activities interplay with each other, and cooperatively regulate life machinery. As reported before, in the crowded cellular environment of biological tissue, motion and shape transitions of a single cell can deliver dynamic mechanical cues (e.g., shearing, compressing, pulling and stretching) to its neighbors^16,17^, which subsequently affect cell signaling cascade. For instance, Schrader et al. revealed that abnormal ECM stiffness causes intracellular structural changes at the cytoskeleton-membrane interface, and thereby block signaling pathways from targeting drugs^6^. Fibroblast-collagen matrix (FCM) can set free soluble cytokines under external mechanical actuations, which activate intra-cellular signaling pathways to regulate cell fate^7^. These results indicate that dynamic mechanical cues resulting from inter-cell interactions can affect the intra-cellular activities. However, a complete study on the interplay between inter- and intra-cellular collective activities, and how they cooperatively regulate macroscopic tissue functions has not yet been conducted.

In this work, we studied the effect of dynamic mechanical cues on the collective behavior of population cells by monitoring the inter- and intra-cellular activities within the confluent cell monolayer (CCM), which is maintained in a customized microfluidic device. Our studies reveal that upon periodic TNF-α stimulation, fibroblasts in CCM show substantially different cellular responses (i.e. elevated NF-κB oscillation frequency and amplitude) as compared to the stand-alone (SA) cells, suggesting the effect of inter-cellular interactions. By mimicking diverse mechanical cues in CCM with the aid of customized microfluidic devices, we demonstrate that the collective behavior within the biological tissue is a result of synergistic actions of macroscopic tissue deformation, mechano-signaling and NF-κB dynamics. Phase-mismatching among those factors leads to disrupted collective behavior. This investigation reveals a novel cascade process linking the inter- and intra-cell collective behavior, in which dynamic mechanical cues and mechano-signaling play crucial roles in regulating the macroscopic function of biological tissues.

## Results

### Inter-cellular Mechanical Cues in CCM upon periodic TNF-α stimulation

To construct a confluent cell monolayer (CCM), 3T3 fibroblasts were cultured at ~100% confluency for 24 h in 96-well plate before being transferred to a shear-free microfluidic device (Fig. S1a-c), where dynamic inflammatory signals of various frequencies and amplitudes are introduced to mimic the everchanging cellular environment *in-vivo* (Fig. S1d-f)^18^. It is demonstrated that fibroblasts self-organize into a monolayer sheet (Fig. S2a-b), and the gap between cells is occupied by collagen fibers (Fig. S2c). When being exposed to periodic TNF-α stimulation, CCM of different sizes exhibit distinctive morphological responses (Fig. S2d and Video S1). When the size is small (i.e. small quantities of cells), CCM exhibits contractile behavior upon periodic TNF-α stimulation, and the morphological transitions are irreversible. In this work, we studied the collective response of cell population containing more than 80 fibroblasts, where the culture chamber of 400 μm by 400 μm in size is fully occupied and the overall size of CCM remain unchanged during periodic TNF-α stimulation.

At the cell-cell and cell-ECM interfaces, mechanical cues including pulling (elongated cell morphology), stretching (increased cell surface area) and compression (decreased cellular volume) are generated by interactions among neighboring cells, and macroscopic tissue deformation (Video S2). Displacements of nuclear centroids show both long-distance migration and vibration with respect to he averaged trajectory (Fig. 1b and Fig. S2e). To evaluate coordination of population cells in CCM, trajectory (*x*_1_, *x*_2_, ⋯ *x*_*n*_) and (*y*_1_, *y*_2_, ⋯ *y*_*n*_), where *n* is the total frame number, is averaged every 5 to 10 steps depending on the frame rate, e.g. 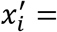 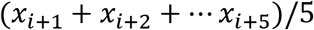 and 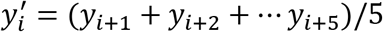 The deviation of cell migration coordinates from the averaged trajectory Δl can then be calculated from its minimum distance to the piecewise cubic interpolation curve of 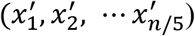 and 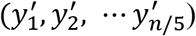 Fig. S2e). We observed that even though the long-distance migration differs among population cells, they shared a similar vibrating frequency, which coincides with the periodicities of TNF-α inputs (Fig. 2d-f), reflecting deformation of CCM as a whole entity.

**Figure 1:**
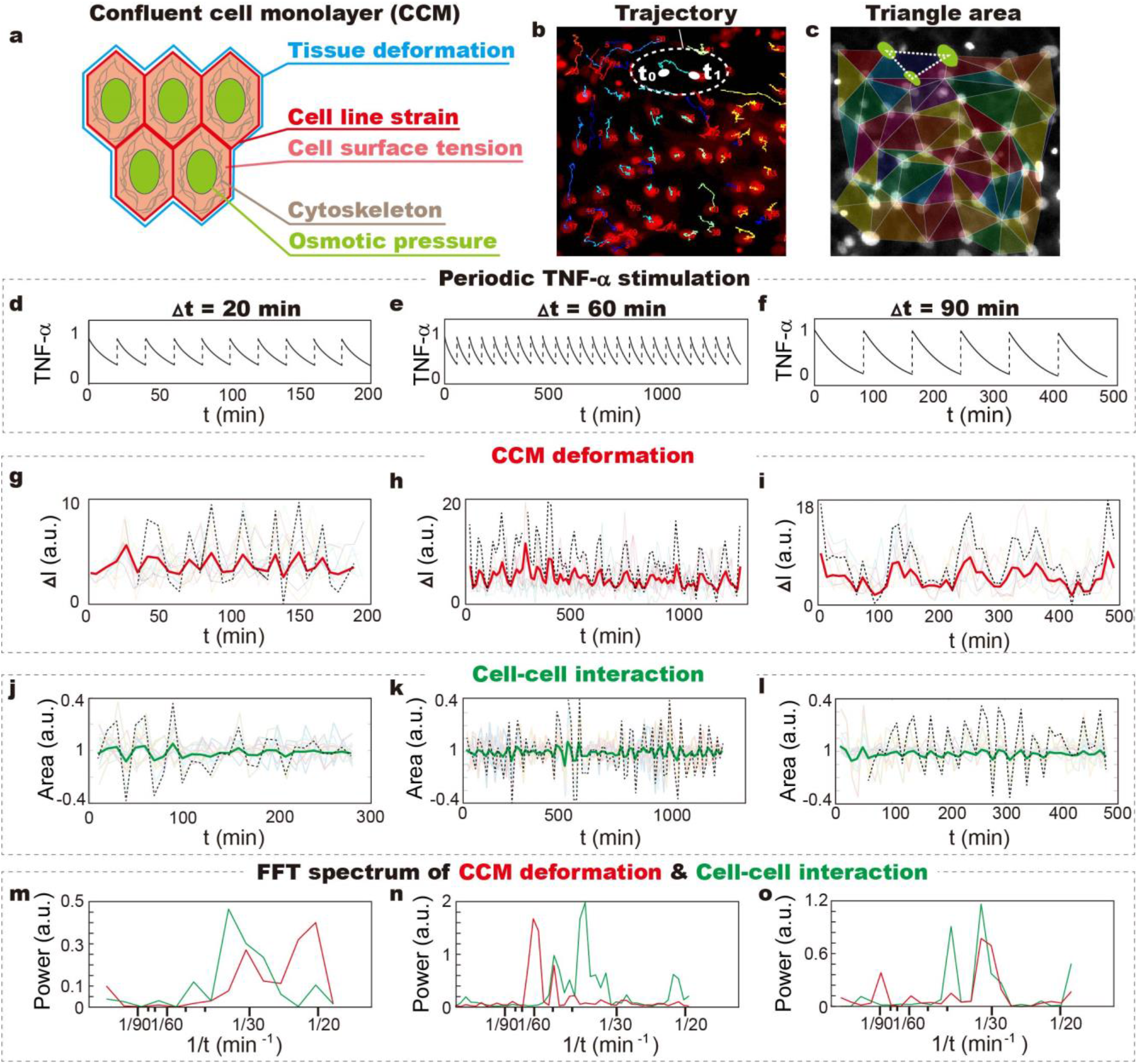
Morphological responses of CCM: periodic TNF-α input induces dynamic mechanical cues in the confluent cell monolayer (CCM). **a.** Schematic shows that various types of mechanical cues can be reflected by morphological features of single cells in CCM, where coordinated vibration of nuclear centroids reflects CCM deformation; cell morphological transition reflects surface tension and line strain; nuclear shape changes reflect intracellular osmotic pressure and cytoskeleton organization. **b.** Trajectories of nuclear centroids upon periodic TNF-α stimulation show distinctive mobilities of single cells. **c.** Cell-cell interactions are evaluated by measuring the triangular area connecting 3 neighboring cells. **d-f.** Periodic TNF-α inputs are generated by controlled delivery of TNF-α into the microfluidic culture chambers. The fluctuation in TNF-α concentration is accomplished by natural protein degradation (Shown in Fig. S1f). **g-i.** Collective vibration of nuclear centroid during cell migration within CCM reflects deformation of cell monolayer, which coordinates with periodic TNF-α stimulation. **j-l.** Relative displacement of nuclear with respect to the neighbors causes changes in the triangles area, which reflect the cell-cell and cell-ECM interactions. **m-o.** Fast Fourier Transform (FFT) shows that variations in the triangular area at all TNF-α input periodicities share a similar dominant frequency, ranging from 1/40 to 1/30 min^−1^. While, the dynamic CCM deformation mostly synchronize with TNF-α input.

**Figure 2:**
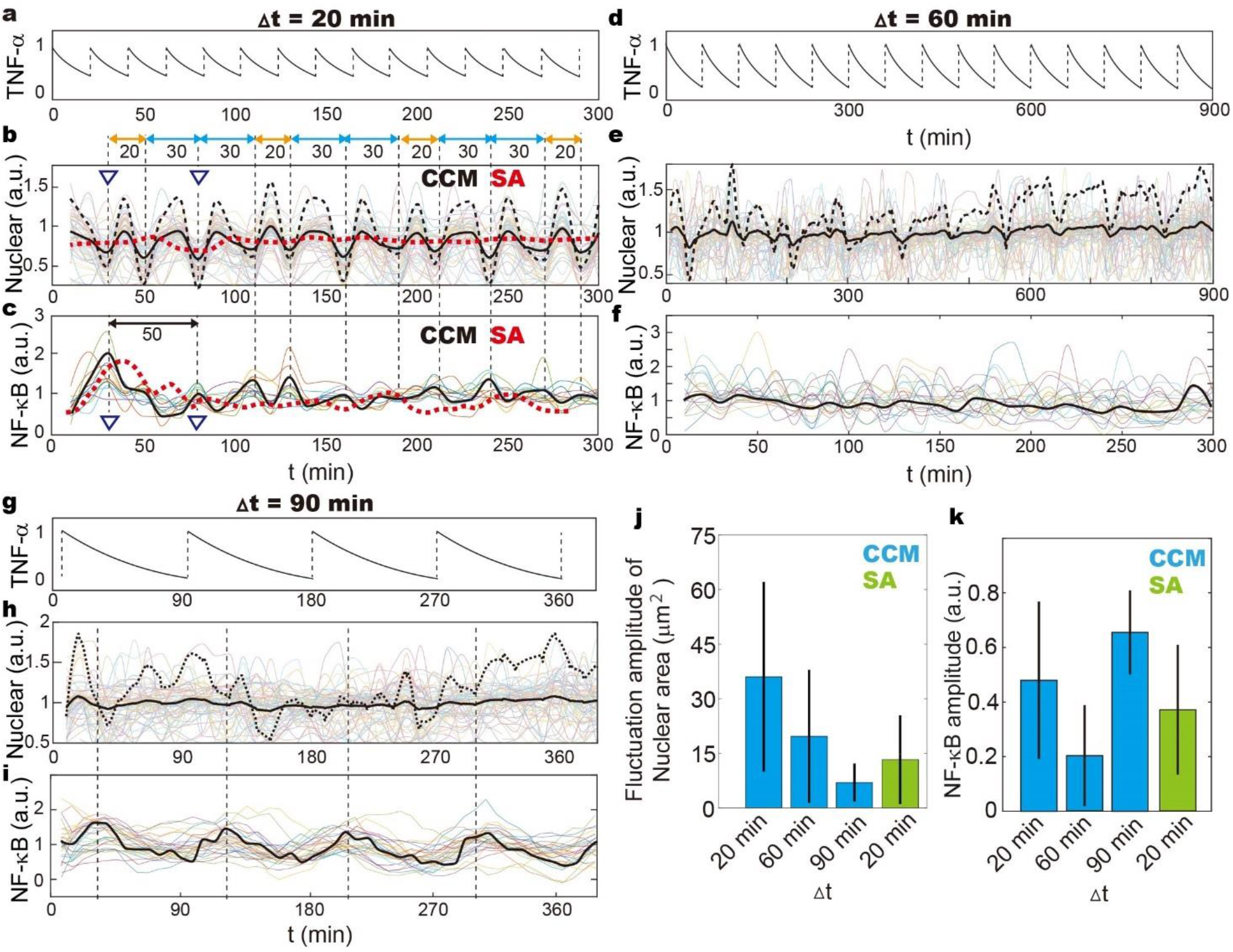
Intracellular collective responses: periodic TNF-α input induces collective activities in nuclear shape fluctuation (NSF) and NF-κB dynamics. NF-κB dynamics is tracked by p65-dsRed and nuclear shape transition represented by H2B-GFP. **a-i.** NF-κB and NSF traces of single fibroblasts in CCM upon stimulation by **a-c.** oscillatory 1/20 min^−1^ TNF-α input, **d-f. 1/** 60 min^−1^ oscillatory TNF-α input, and **g-i. 1/** 90 min^−1^ oscillatory TNF-α input. Solid lines reflect NF-κB dynamics and nuclear area variances averaged among all participating cells. The black dashed lines are the enlarged view of those solid lines, showing subtle variations in NSF, and the red dashed lines in Fig. b and c represent averaged NF-κB trace of stand-alone (SA) fibroblasts. It was observed that 1/20 min^−1^ oscillatory TNF-α input induces collective activities in both NF-κB dynamics and NSF. Even though NF-κB oscillation gets entrained by 1/90 min^−1^ oscillatory TNF-α input, NSF of fibroblasts in FCM is mostly random. **j.** Amplitude of NSF is maximized at 1/20 min^−1^ TNF-α input frequenty, showing substantially higher values than CCM, which is exposed to TNF-α stimulation of other frequencies. **k.** NF-κB oscillation amplitude is maximized at 1/90 min^−1^, when getting entrained, and greatly enhanced in CCM in response to 1/20 min^−1^ TNF-α stimulation as compared to SA cells.

As cells in CCM show distinctive mobilities, the trajectory of nuclear centroids can not reflect mechanical cues at cell-cell and cell-ECM interfaces. To investigate the inter-cellular interactions, we measured the variations of the triangle area connecting 3 neighboring cells (Fig. 1c), which reflect mechanical cues from neighboring cells and in some extent, the intra-cellular cytoskeleton reorganization^19,20^. We observe that collective activities emerge at all TNF-α input frequencies, reflecting synchronized cell-cell interactions (Fig. 1j-i). But, the variations in triangle area do not coordinate with the periodic TNF-α stimulation. Instead, FFT spectrums show dominant frequencies between 1/30 min^−1^ and 1/40 min^−1^ regardless of TNF-α input periodicities (Fig. 1m-o), suggesting the frequency is a character intrinsic to CCM. In contrast, the morphological responses of single fibroblasts (i.e. outline of the cell reflecting line strain, cell area reflecting surface tension,) are highly heterogenous and random with dynamic TNF-α stimulations (Fig. S2f-k).

### Intra-cellular Collective behavior upon periodic TNF-α stimulation

The intra-cellular mechanical cues are assessed by monitoring changes in the nuclear morphological features, which reflect variations in osmotic pressure and cytoskeleton reorganization (Fig. 1a)^21^. Our results demonstrate that collective behavior emerges only at 1/20 min^−1^ TNF-α input periodicity, showing synchronized decrease in nuclear area (Fig. 2b, 2e, 2h). We observe that cell nucleus quickly deforms showing increased H2B fluorescence intensity, which suggests more densely packed chromatin fiber, and restore its original shape within 20 to 30 min (Fig. S2a). Notably, the fluctuation in nuclear shape does not synchronize with TNF-α input, showing mode hopping between 1/20 min^−1^ and 1/30 min^−1^ (Fig. 2b). The amplitude of nuclear shape fluctuation (NSF) is also considerably higher at 1/20 min^−1^ as compared to 1/60 min^−1^ and 1/90 min^−1^ TNF-α inputs (Fig. 2j). The deviation of nuclear shape fluctuation (NSF) frequency from TNF-α input periodicity and dynamic mechanical cues in CCM suggests the participation of multiple cycling processes.

Besides the collective morphological responses, NF-κB dynamics of population cells get synchronized as well at 1/20 min^−1^ TNF-α input frequency (Fig. 2c). Cells in CCM shows apparently stronger response than the stand-alone (SA) cells, which is reflected by elevated fluctuation amplitude of nuclear NF-κB fluorescence intensity (Fig. 2c) and the increased number of activated cells (Fig. S2k). For example, the fraction of active cells is ~ 20% higher in CCM than the isolated ones when the input TNF-α concentration (indicated by dashed line) is 0.08 ng/ml. Notably, with 1/20 min^−1^ TNF-α stimulation, the NF-κB dynamics shows an oscillating frequency of ~ 1/26 min^−1^, which coordinates with the collective action of NSF, and considerably more frequent than its 1/90 min^−1^ natural value^15^. As fibroblasts in CCM are genetically identical with the SA cells, the stronger and coordinated NF-κB activities of fibroblasts can be attributed to the dynamic mechanical cues at multiple scales including the macroscopic CCM, the inter- and the intra-cellular level.

### Mechanical cues at cell-ECM interface is insufficient to enhance NF-κB dynamics

To eliminate the environmental complexities, we systematically studied the effect of different types of mechanical cues on NF-κB dynamics with the aid of microfluidic devices. To mimic the inter-cell interactions in CCM, we firstly loaded the mixture of fibroblasts and collagen solution into the microfluidic chip. At 37℃, collagen solution quickly solidifies and forms cell-ECM interface surrounding SA cells. As cell movement causes uncontrollable mechanical cues delivering to its neighbors, fibroblasts are examined in the microfluidic device at the single cell level (Video S3). The programmed on-off of 4 control lines, which are connected to stretchable PDMS membrane of 2 mm in diameter, causes collagen matrix remodeling and thus induces dynamic mechanical cues to SA cells (Fig. 3a). The amplitude and types of induced morphological fluctuation (IMR) depends on the inflating pressure as well as the location of the cells with respect to the PDMS membrane, i.e. pulling (elongated cell morphology), stretching (increased cell surface area) and compression (decreased cellular volume) (Fig. S4c-e). TNF-α inputs of 1/20, 1/60 and 1/90 min^−1^ periodicities and dynamic collagen remodeling of 1/20 to 1/90 min^−1^ frequencies are then simultaneously introduced to SA cells. We demonstrate that NF-κB dynamics is only slightly enhanced when SA cells are compressed, and when dynamic mechanical cues are at 1/20 min^−1^ frequency or mode hopping between 1/20 and 1/30 min^−1^ (Fig. 3b and Fig. S4f-g). Even though NF-κB oscillation frequency of few SA cells reach 1/30 min^−1^, the enhanced and coordinated NF-κB dynamics comparable to CCM is not yet accomplished. In the meantime, the induced mechanical cues at cell-ECM interface bring no observable changes in NSF amplitude at all TNF-α input frequencies. The failing in reconstituting the intracellular morphological and signaling responses in CCM suggests that NF-κB dynamics is closely associated with NSF, and possibly the mechano-signaling, activation of which leads to cytoskeleton reorganization and consequently induces nuclear shape changes.

**Figure 3:**
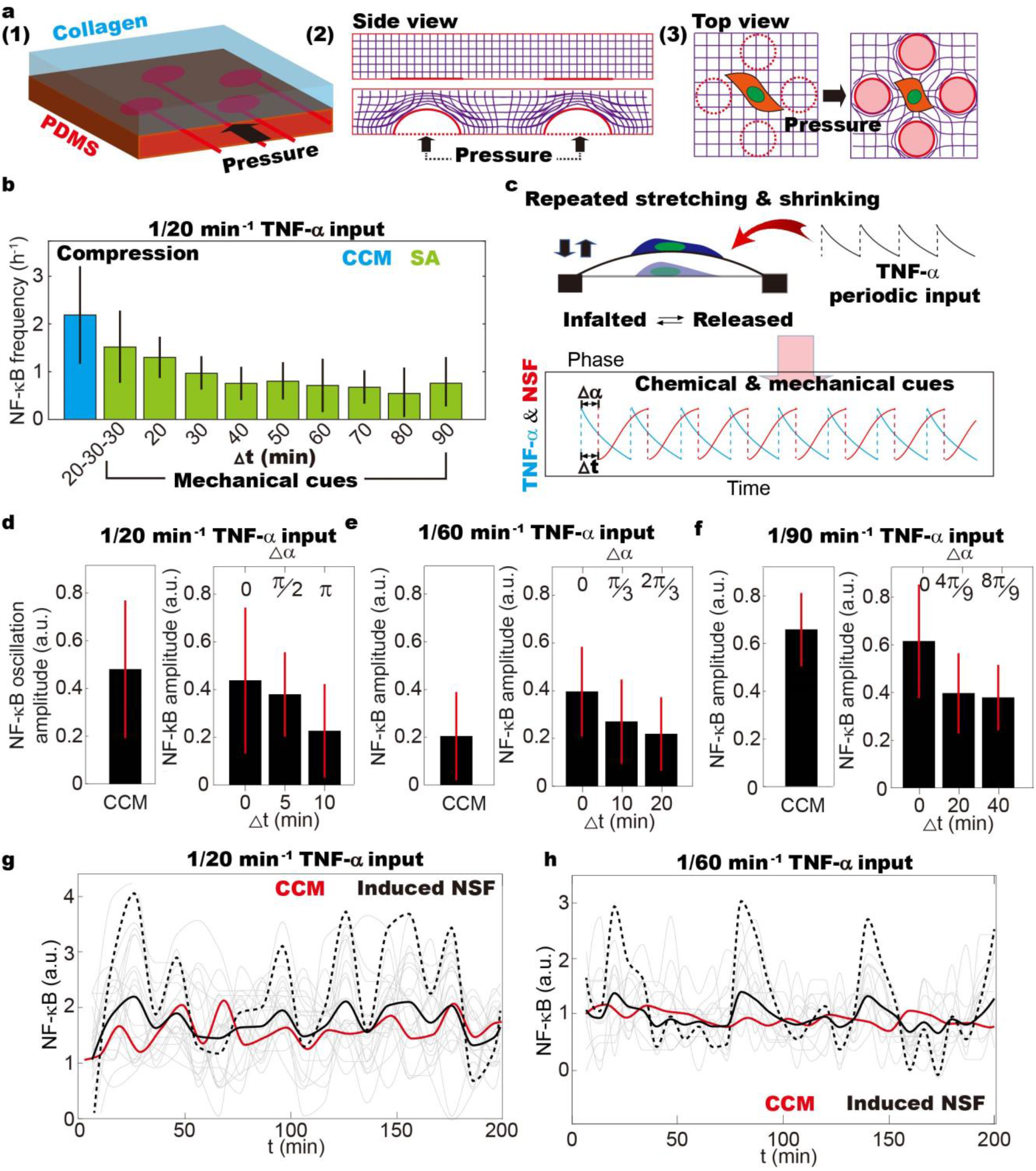
Modeling the inter- and intra-cellular mechanical cues in CCM: the effect of dynamic mechanical cues on NF-κB oscillation of SA cells. **a.** Schematic drawing shows that the organization of collagen fibers can be dynamically altered by repeatedly pressurizing and relaxation of the underneath PDMS membrane. The remodeling of collagen fiber matrix then delivers dynamic mechanical cues to SA cells. **b.** By simultaneously introducing 1/20 min^−1^ TNF-α and different frequencies of collagen remodeling ranging from 1/20 to 1/90 min^−1^, the effect of CCM contraction on NF-κB dynamics is systematically studied. It is observed that when cell are compressed, the induced mechanical cues are insufficient to achieve NF-κB oscillation frequency comparable to CCM. But, NF-κB dynamics is enhanced, when the frequency of induced collagen remodeling is 1/20 min^−1^ or mode hopping between 1/20 and 1/30 min^−1^. **c.** NSF and TNF-α inputs are simultaneously introduced to SA cells using customized microfluidic chip. The time (Δ*t*) and phase (Δα) difference between minima of the nuclear area (red) and maxima of TNF-α concentration models the phase-mismatching between mechanical and chemical cues in CCM. **d-f.** Synergy between NSF and TNF-α inputs induce elevated NF-κB oscillation amplitude among SA cells, which is comparable to or even higher than the ones in CCM. It is suggested that CCM as a resonant system, may have an intrinsic frequency of ~ 1/20 min^−1^, deviation from which leads to diminished NF-κB amplitude. **g-h.** Synergy between NSF and TNF-α inputs induces collective NF-κB oscillation activities coordinating with TNF-α periodicity. NF-κB dynamics averaged among all cells with induced NSF (black lines) shows higher amplitude as compared to the ones in CCM (red lines) when being stimulated by 1/60 min^−1^ oscillatory TNF-α input. The values are comparable in CCM and on-chip in response to 1/20 min^−1^ oscillatory TNF-α input. The dashed lines are the enlarged view of the black lines.

### Dynamic nuclear shape fluctuation facilitates NF-κB dynamics

To verify our hypothesis, nuclear shape fluctuation (NSF) is induced on SA cell by directly plating fibroblasts on the stretchable PDMS membrane (Fig. S5a). Stretching or shrinking of cell adhering PDMS surface alters cell adhesion area, and consequently induce NSF mimicking the cellular behavior in CCM (Fig. S5a-c)^22,23^. We demonstrate that the induced NSF is morphologically similar to CCM upon TNF-α stimulation. For example, if cells are initially plated on the inflated membrane at t+0 min (i.e. initially), the adhesion area will be suddenly decreased when the pressure is released at t+10 min (Fig. S5c). SA cells are compressed at this moment with maximized nuclear height, and gradually restore their original conformation within ~10 min (Fig. S5b-c). Intriguingly, we observe that the responsiveness of fibroblasts changes during induced NSF (Fig. S5e). SA cells are more responsive to TNF-α stimulation when being compressed (Fig. S5b), and the fraction of active cells decreases as cells go back to its original shape (Fig. S5e). Studies by Agnès, et al. reveal that the signaling cascades can be regulated by variances in cell and nuclear shape, which causes cytoskeleton reorganization and unbalances osmotic pressure across NE^4,24^. Apparently, when the collective NSF emerges at 1/20 min^−1^ TNF-α input frequency, population cells in CCM are repeatedly compressed (i.e. decrease in nuclear area shown in Fig. 2b), and forced to a more responsive state (Fig. S5f). Therefore, NF-κB shuttling coordinates with NSF, and shows collective oscillation behavior more frequent than its intrinsic value (Fig. 2c).

We then repeatedly stretch or shrink the cell adhesion surface and simultaneously introduce periodic TNF-α stimulation to SA cells (Fig. 3c). It is demonstrated that during 1/20 min^−1^ periodic TNF-α stimulation, phase-matching between NSF and TNF-α inputs (Δt = 0, Δφ = 0) leads to increased NF-κB oscillation amplitude comparable to the cellular responses in CCM, and twice the value of the control cells (Fig. 3d). Enlarged phase difference (Δt = 5 and 10 min; 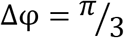 and 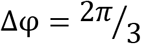) causes decreased nuclear NF-κB level. In the meantime, the otherwise random NF-κB dynamics of SA cells as shown in Fig. S2g, get more coordinated with synergized NSF and TNF-α input, showing shared frequency of ~ 1/20 min^−1^ (Fig. 3g)^25^. Similar effect was observed with 1/60 min^−1^ and 1/90 min^−1^ TNF-α inputs as well (Fig. 3e-f and 3h). We thus conclude that the collective NSF activities in CCM facilitate NF-κB dynamics, in which the phage-matching of dynamic chemical inputs and NSF is crucial. Intriguingly, NF-κB dynamics gets coordinated and enlarged (i.e. entrainment) upon periodic TNF-α stimulation at its natural frequency of 90 min (Fig. 2i, 2k and Fig. 3f)^15,25^. The collective cellular responses can be disrupted by phage-mismatching mechanical cues inducing NSF, which further emphasizes the importance of synergy between intracellular mechanical cues and the chemical signals (Fig. 3f).

### Mechano-signaling causes nuclear shape fluctuation

So far, we have demonstrated that periodic TNF-α inputs causes dynamic mechanical cues propagating in CCM, and determined the cause and consequence relationship between intracellular collective morphological and signaling responses. But, the correlation between the dynamic mechanical cues and NSF remains unclear. It is worth noting that the emergence of collective NSF activities at a defined TNF-α input frequency is similar to the entrainment in the signaling cascade^4^, which is brough up by Tay, etc. to interpret coordinated and enlarged signaling activities. Meanwhile, previous studies reveal that activation of mechano-signaling pathways leads to cytoskeleton reorganization, which subsequently induces nuclear shape changes^21,26^. We, therefore, hypothesize that mechano-signaling is triggered by the dynamic mechanical cues in CCM, which actively regulate NSF, and consequently facilitate NF-κB dynamics.

To study the role of mechano-signaling, both CCM and the SA cells are treated with the Rho-associated protein kinase (ROCK) inhibitor Y-27632, which plays central roles in multiple mechano-transduction pathways^27^ and is closed associated with cytoskeleton reorganization^28^. Our results demonstrate that with the application of ROCK inhibitor, CCM collective activities in both NSF and NF-κB dynamics at 1/20 min^−1^ TNF-α input frequency are disrupted (Fig. 4a and 4b). In the meantime, no distinguishable transition is observed in CCM at 1/60 and 1/90 min^−1^ TNF-α input frequencies, and SA cells under all TNF-α stimulation conditions (Fig. 4b). Notably, the enhancement of NF-κB dynamics, which is brought up by induced NSF, is also not affected (Fig. 4b). These results suggest that mechano-signaling is a crucial link between external mechanical cues and collective intra-cellular behaviors, which causes synchronized NSF, and subsequently leads to elevated NF-κB activities at 1/20 min^−1^ TNF-α input frequency. The link can, however, be omitted if NSF synergizing with the periodic TNF-α input can be induced to CCM and SA cells.

**Figure 4:**
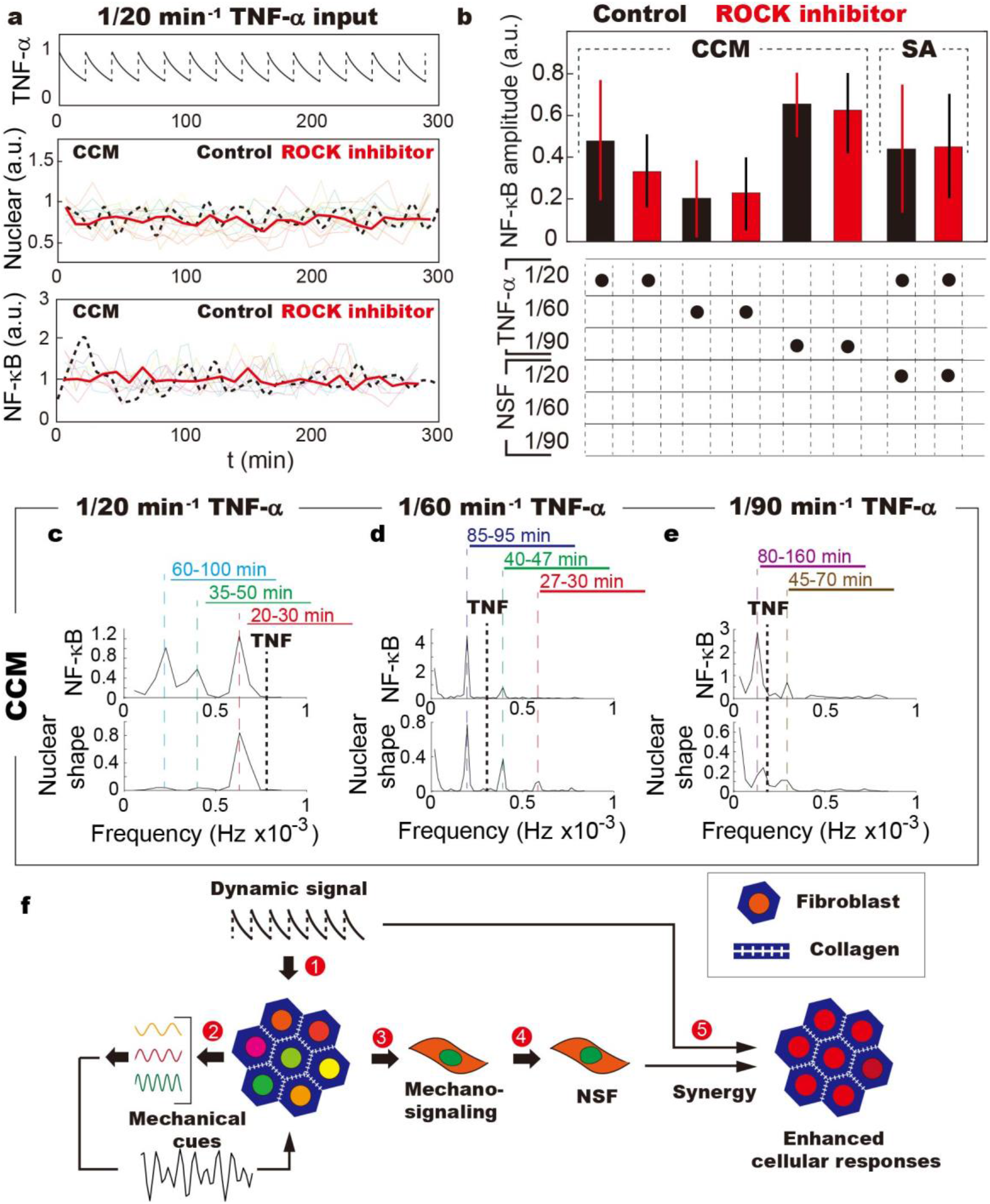
The role of mechano-signaling: ROCK inhibitor Y-27632 inhibit CCM collective activities in NSF and NF-κB dynamics. **a.**NF-κB and NSF traces of single fibroblasts in CCM treated with ROCK inhibitor, which is followed by 1/20 min^−1^ TNF-α stimulation. It is demonstrated that the application of ROCK inhibitor disrupts the collective activities of CCM in NSF and NF-κB dynamics shown in Fig. 1b-c. **b.** The oscillation amplitude of NF-κB in CCM is affected only at 1/20 min^−1^ TNF-α input frequency, indicating mechano-signaling is not activated under other conditions (i.e. 1/60 and 1/90 min^−1^). The lack of transitions with induced NSF on SA cells suggests NSF is caused by downstream reactions of mechano-signaling pathways. **c-e.** Fast Fourier Transform (FFT) shows dominant NF-κB oscillation and NSF frequencies of fibroblasts in CCM upon stimulation by 1/20 min^−1^ (c), 1/60 min^−1^ (d), and 1/90 min^−1^ (e) periodic TNF-α stimulations. It is demonstrated that in CCM, NF-κB oscillation closely correlates with NSF, showing similar dominant frequencies, (i.e. ~ 1/26 min^−1^ with 1/20 min^−1^ TNF-α stimulation; ~ 1/90 min^−1^ for 60 min TNF-α stimulation; and ~ 1/100 min^−1^ with 90 min^−1^ TNF-α stimulation). **f.** Schematic showing that collective activities of population cells in CCM is initiated by mechano-signaling. ① When periodic chemical inputs meet the natural frequency of CCM, morphological responses of population cells generate various types of mechanical cues; ② The coupling of these mechanical cues leads to complex signal (e.g. mode hopping between 1/20 and 1/30 min^−1^); ③④ Activation of mechano-signaling leads to cytoskeleton reorganization, and thus NSF; ⑤ Synergy between NSF and TNF-α input leads to enhanced cellular responses (i.e. elevated NF-κB oscillation frequency and amplitude).

## Discussions

This study reveals that in response to dynamic stimuli, mechanical cues propagating within the biological tissues facilitate collective cellular responses, including nuclear shape fluctuation (NSF) and intra-cellular signaling activities. Notably, only at a specific TNF-α input frequency of 1/20 min^−1^, mechano-signaling is activated, which subsequently causes elevated NSF and NF-κB dynamics. To the best of our knowledge, it remains puzzling that how the mechano-signaling pathways respond to dynamic mechanical cues. We suspect that similar to the entrainment in biochemical signaling pathways^4^, collective NSF of population cells emerges when the characteristic frequencies of dynamic mechanical cue meet the cycling time of mechano-signaling cascade. Consistently, NF-κB oscillation frequency in response to 1/20 min^−1^ TNF-α stimulation is ~ 1.6 times higher when the periodicity of the induced mechanical cues is close to NSF in CCM. The failing in causing NSF with and without TNF-α stimulation suggests that the activation of mechano-signaling pathways may require coupling of multiple dynamic mechanical cues, and even intra-cellular biochemical reactions related to TNF-α stimulation. The random fluctuation in single cell line-strain and surface tension may also contribute as essential noise (Fig. 4f)^4^.

The cause and consequence relationship between NSF and elevated NF-κB dynamics is determined by simultaneously inducing dynamic NSF and periodic TNF-α stimulation. Previous studies have shown that variations in nuclear shape can unbalance the osmotic pressure across NE and affects transportation of nuclear proteins^29^. Similar phenomena were observed by inducing SA cells nuclear shape changes on a stretchable PDMS membrane (Fig. S3j-k). Fibroblasts are indeed more responsive when the nuclear shape experiences similar decrease as the cells in CCM. It is plausible that the cycling time of involved mechano-signal pathways, which leads to cytoskeleton reorganization and nuclear shape changes, is considerably less than NF-κB dynamics. At 1/20 min^−1^ TNF-α frequency, coordinated NSF of population cells collectively forces fibroblasts to a more responsive state, and consequently expedite the NF-κB oscillation frequency. The hypothesis is consistent with our observation that NF-κB activities of different SA cells synchronize as long as TNF-α input phage-matches with NSF (Fig. 3d-h). The positive effect of NSF on NF-κB dynamics is further verified by Fast Fourier Transform (FFT) analysis of NSF and NF-κB dynamics. The high frequency NF-κB oscillations, which approximately locate at 1/26 min^−1^, only emerges when the dominant frequency of NSF is at the same position (Fig. 4c-d).

Unlike the common belief that the effect of physical environmental conditions is long-term^30–32^, our studies reveal that dynamic mechanical cues originating from inter-cell interactions can actively facilitate collective cell responses. In response to dynamic environmental stimulation, synergy between intracellular mechanical cues and extracellular signals leads to enhanced biochemical signaling cascade, i.e. cycling frequency and amplitude^33–45^. We postulate that besides its biological significance, our studies may promote the development of therapeutic approaches, in which the application of dynamic mechanical cues to patients though acoustic forces or magnetic field facilitates drug delivery and activates local immune responses.

## Supporting information

Supplementary Vdieo 1

Supplementary Vdieo 2

Supplementary Vdieo 3

