## Supplementary Information for "Dynamic Mechanical Cue Facilitate Collective Responses of Crowded Cell Population"

### Movie descriptions

**Supplementary Video 1:** Collective movement of fibroblasts when being maintained in CCM. The connective sheet partially detached from surface, showing contractile actions.

**Supplementary Video 2:** Morphological responses of fibroblasts in CCM when being stimulated by (from left to right) 1/20 $min^{-1}$, 1/60 $min^{-1}$, 1/90 $min^{-1}$ oscillatory TNF-α input and single pulse TNF-α stimulation. Traces of single cell nuclear area (H2B-GFP) variations over time are plotted in the lower panels

**Supplementary Video 3:** Movement of fibroblasts in the collagen matrix suggests that actions of single cells can deliver mechanical cues to the neighbors. It is also demonstrated that it takes longer time for SA cells to attached to the collagen fiber as compared to the fibronectin-coated substrates.

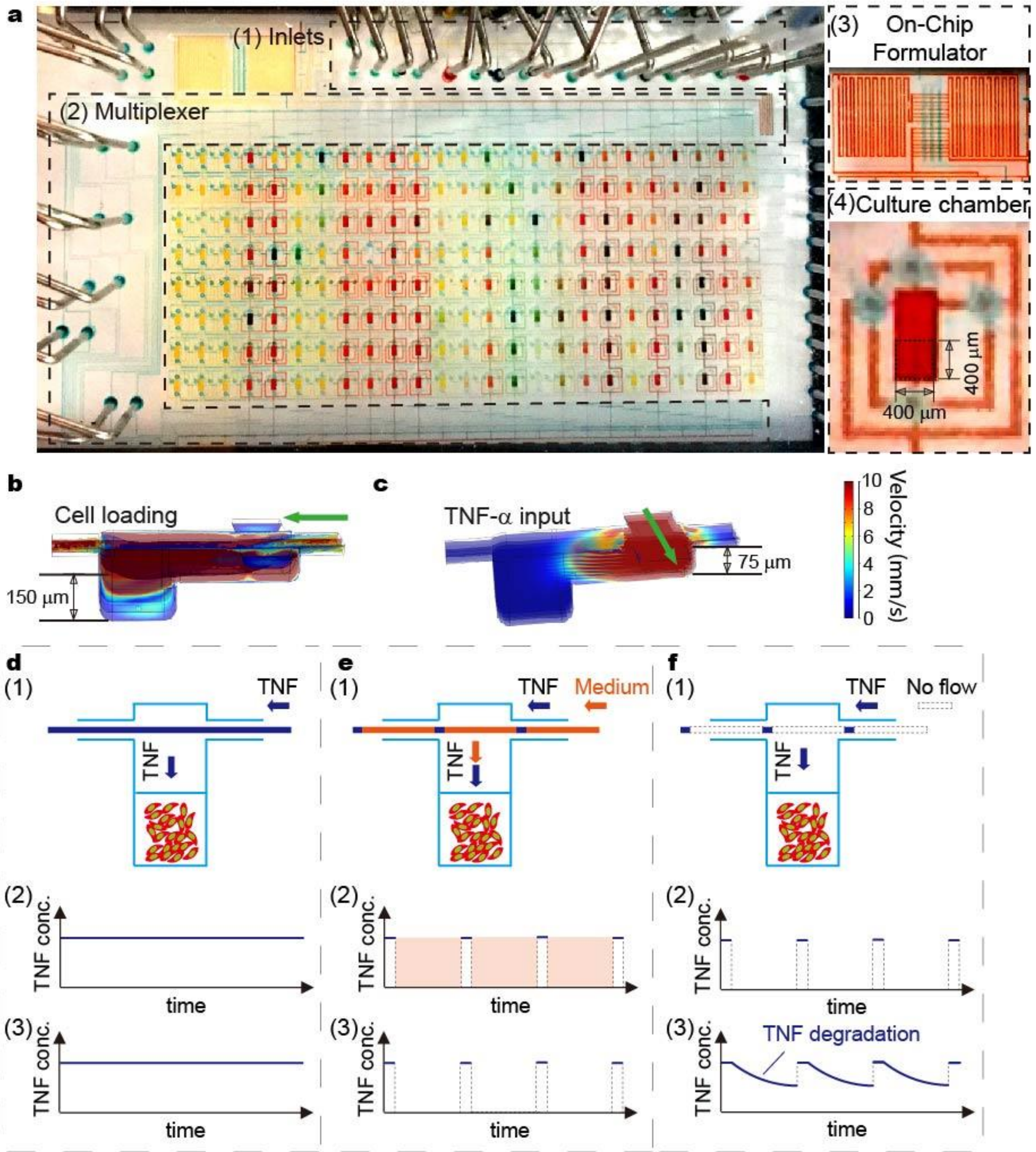


**Supplementary Fig.1 Fibroblasts in CCM and stand-alone (SA) cells are maintained in a shear-free microfluidic device. a,** Automated cell culture system for dynamical live-cell analysis. (1) The microfluidic device contains 15 inlets; (2) a multiplexer, which allows control of 200 culture units; (3) an on-chip formulator with nanoliter accuracy; (4) and shear-free culture units controlled by 4 active valves (blue). The culture units are composed by 2 parts, a culture chamber of $400\ \mu\text{m} \times 400\ \mu\text{m} \times 150\ \mu\text{m}$ in dimension and a buffering level of $400\ \mu\text{m} \times 400\ \mu\text{m} \times 75\ \mu\text{m}$. **b,** Two-layer cell culture chamber allows firstly quick cell loading. **c,** Nutrients and TNF-α are delivered to cells in a shear-free manner. **d-f:** Dynamic inflammatory signals generated using microfluidic module: **d,** Continuous delivery of fresh TNF-α solution to cultured cells in diffusion-based mode maintain TNF-α concentration in cellular microenvironment. **e,** Timely replacement of TNF-α solution by culture medium allows generation of pulsatile TNF-α input. **f,** Programed delivery of TNF-α solution generated damped oscillatory TNF-α signal due to TNF-α degradation.

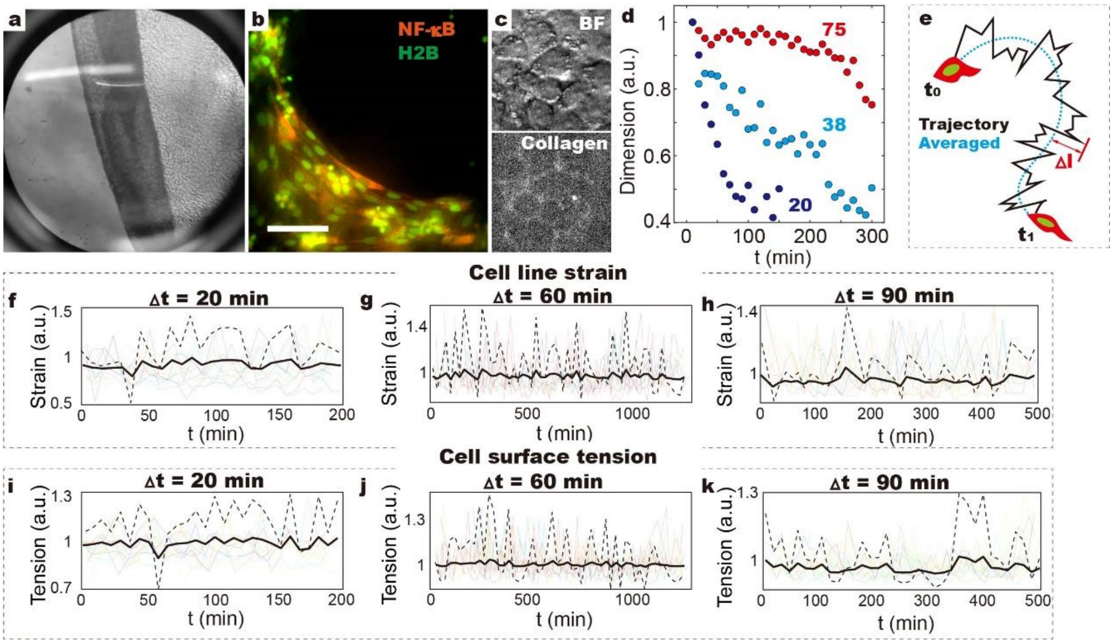


**Supplementary Fig.2 Morphological responses of fibroblasts in the confluent cell monolayer (CCM). a.** Folded CCM layer in 96-well plate shows that fibroblasts form a connective sheet**. b.** CCM layer maintained in the shear-free microfluidic chip (see also Supplementary Video 1). **c.** Bright field (BF) and fluorescent images showing that the gaps between neighboring cells are filled by collagen fibers. **d.** The contractile behavior depends on the size of CCM. For example, CCM containing ~20 cells shrinks down to ~40% of its original size. **e.** CCM deformation is evaluated by measuring the distance of cell migration trajectory to the averaged curve, fluctuation of which reflect vibration of nuclear centroids. **f-h.** Measuring the perimeter of single cells in CCM reveal the line strain, which shows no collective behavior**. i-k**. The variations in 2D area of single cell surface reflect the fluctuating surface tension. Scale bar denote 20 μm.

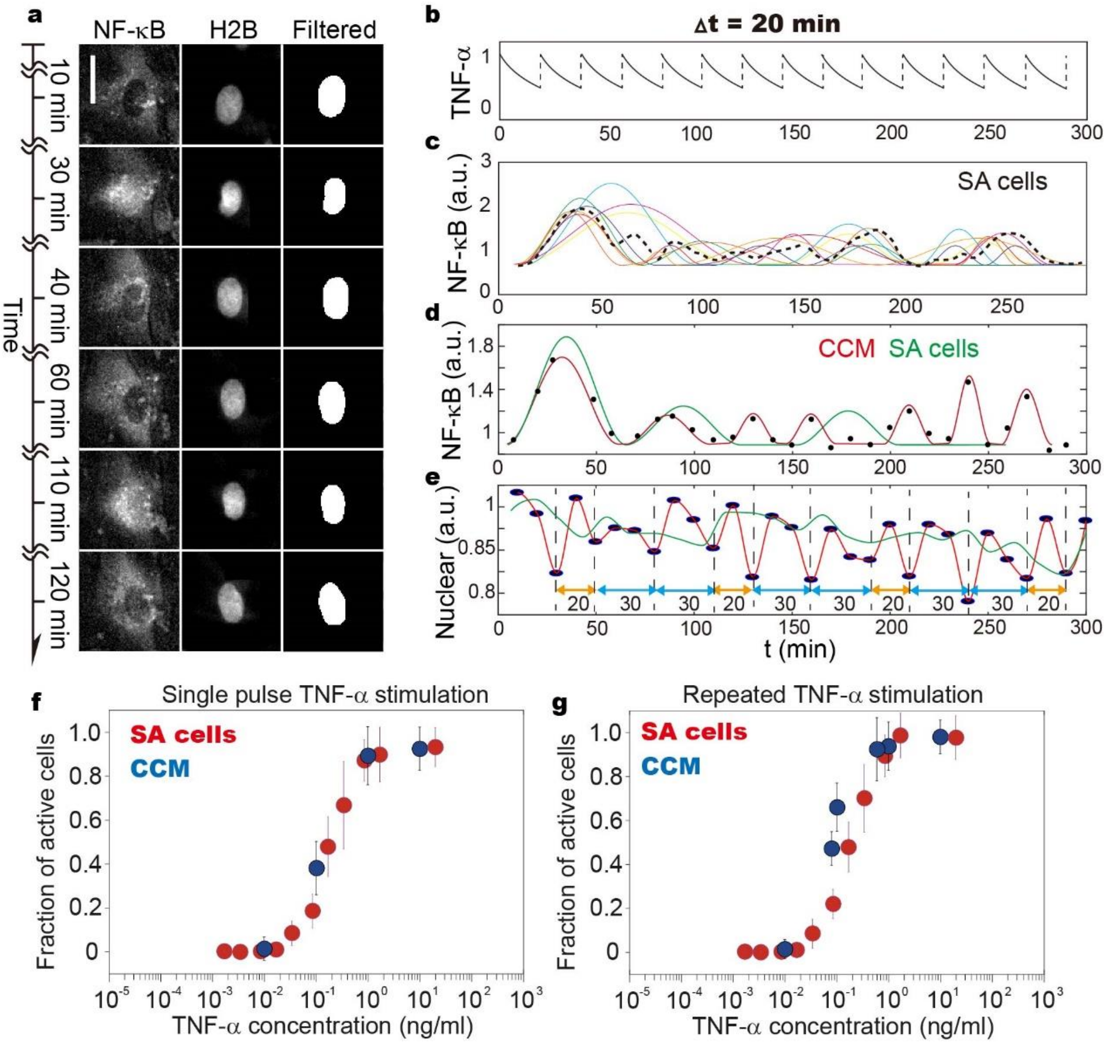


**Supplementary Fig.3 Responsiveness of fibroblasts in CCM and the SA cells. a.** NF-κB dynamics and NSF of single cells in CCM are monitored in real-time using fluorescence microscopy. Images show NF-κB (p65-dsRed) nuclear localization and nuclear shape changes upon periodic TNF-α stimulation. The images of H2B-GFP are processed using real-space bandpass filters to extract morphological features including perimeter and area, etc. **b-e.** NF-κB dynamics and NSF traces of CCM and SA cells when being exposed to 1/20 $min^{-1}$ TNF-α stimulation. It is demonstrated collective activities only emerge in CCM, where amplitude of NSF and NF-κB oscillation is greatly elevated. **f-g.** Fraction of cells responding to (j) single pulse, and (k) repeated TNF-α stimulation (1/20 $min^{-1}$ frequency). It is demonstrated that similar number of fibroblasts in CCM (blue) respond to single pulse TNF-α stimulation as the SA cells (red). While, higher number of responsive cells was observed in CCM as compared to SA cells upon stimulation by repeated TNF-α stimulation.

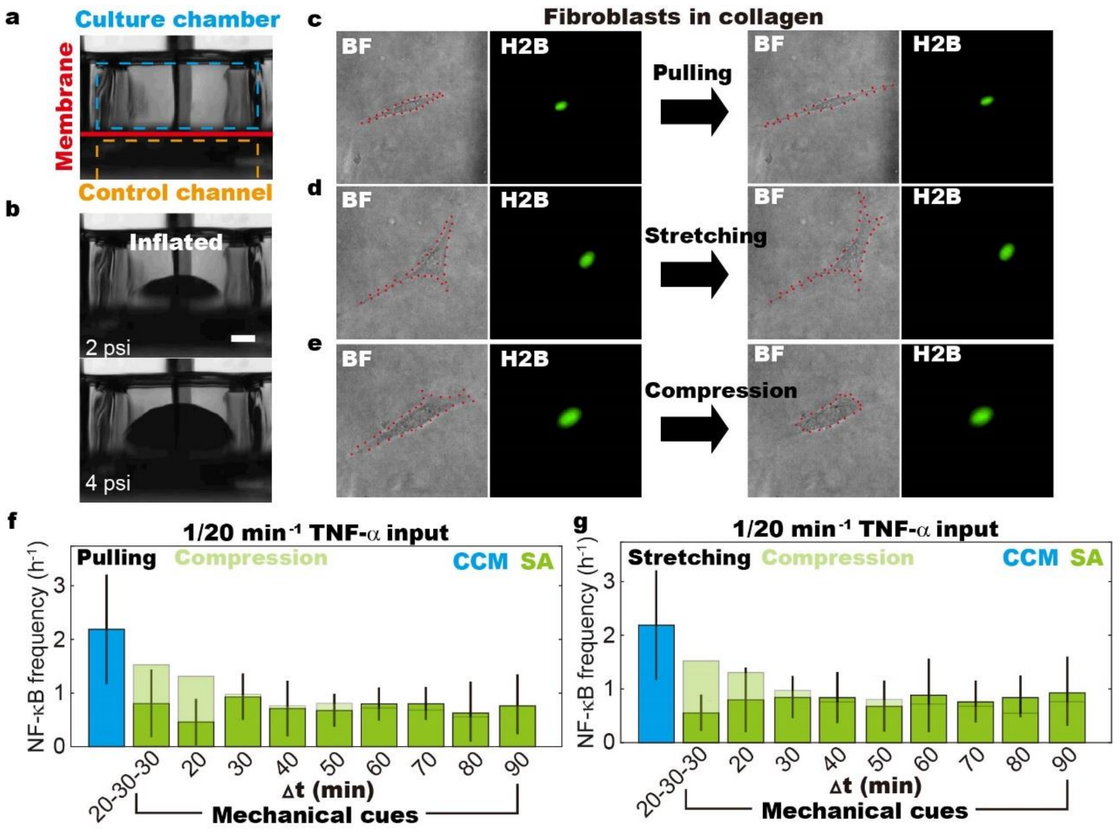


**Supplementary Fig.4 Morphological transitions of stand-alone (AS) cells induced by pressurizing the PDMS membrane, and consequently the remodeling of collagen fibers. a.** Cross-sectional view of the fluidic chip with stretchable PDMS membrane. Fibroblasts are loaded, and maintained in the culture chamber (blue). PDMS membrane (red) is stretched by pressurizing the control channel (orange). **b.** Stretched PDMS membrane at pressure of 2 psi (left) and 4 psi (right). Scale bar denotes 200 μm. **c-e.** Mechanical cues including pulling, stretching and compression are delivered to SA cells by inducing remodeling of collagen fibers. **f-g.** By simultaneously introducing 1/20 $min^{-1}$ TNF-α and different frequencies of collagen remodeling ranging from 1/20 to 1/90 $min^{-1}$, the effect of CCM contraction on NF-κB dynamics is systematically studied. It is revealed that when cells are either pulled or stretched, the induced mechanical cues bring no obvious changes as compared to the SA cells with no induced morphological transitions.

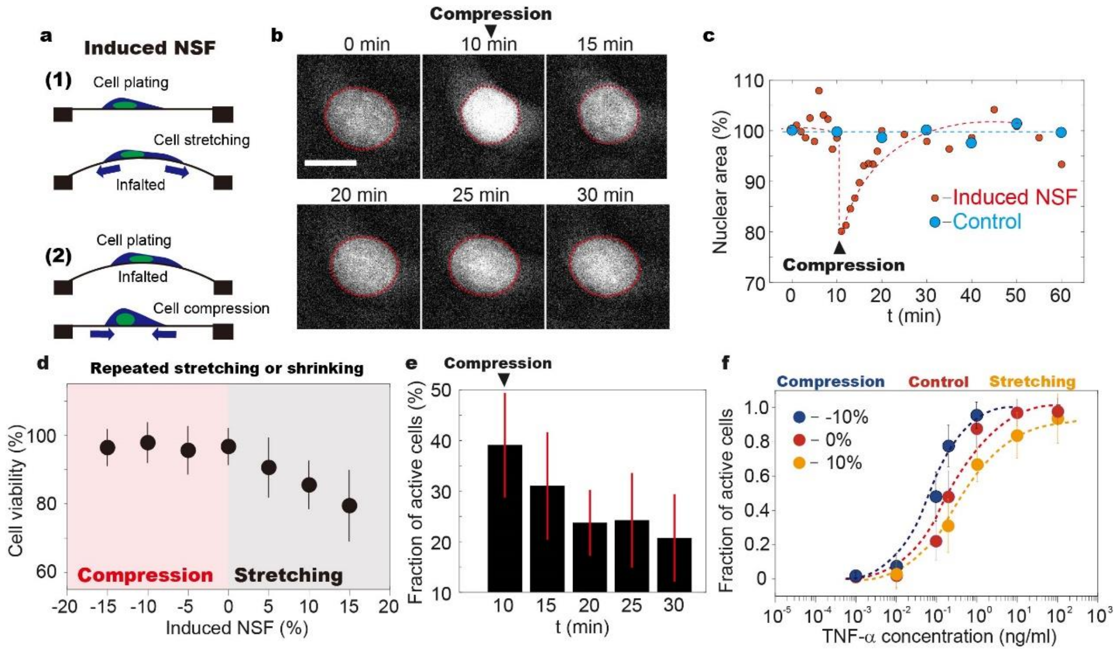


**Supplementary Fig.5 Responsiveness of SA cells with induced mechanical cues and NSF. a.** Schematic view showing that (1) cell adhesion surface can be stretched when being plated on relaxed PDMS membrane, which is later inflated; (2) When cells are plated on stretched PDMS surface, the adhesion area can be decreased by pressure unloading. **b.** Representative fluorescence images of nuclear shrinkage with decreased adhesion area. Cell gradually restore its original conformation within 20 min. Scale bar denotes 10 μm. **c.** Single cell traces showing nuclear shape variances with decreased (red) and unchanged (blue) adhesion area. **d.** Cell viability when being placed on repeatedly stretched or shrunk (at frequency of 1/20 $min^{-1}$) PDMS membrane for 2 hours. It is demonstrated that fibroblasts viability remains similar to control (~100%) when cell adhesion area repeatedly decreases. Cell viability decreases when being stretched, suggesting DNA damage induced by increased osmotic pressure at the top of chromatin. **e.** Responsiveness of fibroblasts to 0.1 ng/ml TNF-α stimulation at different stages of induced nuclear shape change. It is demonstrated that the increased responsiveness upon nuclear shape changes is only temporal. Fibroblasts restore the characteristics of isolated cells after 5 to 10 min. **f.** Fibroblasts are less responsive to TNF-α stimulation (single pulse) when nuclear area laterally expands by ~10%, and more when nuclear area decreases (~-10%).
